# Variations in cellular unfolded protein response, respiratory capacity, and stress tolerance in skin and lung fibroblasts of deer mice (*Peromyscus maniculatus*)

**DOI:** 10.1101/2023.12.07.570632

**Authors:** Kang Nian Yap, KayLene Yamada, Shelby L Zikeli, Yufeng Zhang, Youwen Zhang, Asieh Naderi, Elham Soltanmohammadi, Andreas N Kavazis, Michael D Roberts, Hippokratis Kiaris, Wendy R Hood

## Abstract

Evolutionary physiologists have long been interested in physiological mechanisms underpinning variation in life-history performance. Recent efforts to elucidate these mechanisms focused on bioenergetics and oxidative stress. One underappreciated area that could play a role in mediating variation in performance is the unfolded protein response (UPR), a cellular stress response that reduces secretory protein load, enhances endoplasmic reticulum (ER) protein folding and clearance capacity during stress and during its adaptive phase. Given that the ER and mitochondria interact to regulate cellular homeostasis, it seems intuitive that UPR phenotype would correlate strongly with mitochondrial physiology, which in turn would contribute to variations in whole-organism metabolism. One way researchers have been studying cellular controls of life-history traits is by assessing stress resistance and bioenergetic properties of primary dermal fibroblasts. However, it is unclear if findings from dermal fibroblasts can be generalized to other cell and tissue types, and if fibroblasts’ phenotypes are repeatable across different life-history stages. This study aimed to explore the relationships between UPR profile, cellular respiration, and stress resistance using primary dermal fibroblasts isolated at puberty and primary lung fibroblasts isolated at adulthood. Specifically, we tested if 1) UPR profile of dermal fibroblasts isolated at puberty corresponds to UPR profile of lung fibroblasts isolated at adulthood, 2) UPR profile of dermal fibroblasts isolated at puberty and lung fibroblasts isolated at adulthood correspond to cellular bioenergetics of lung fibroblasts isolated at adulthood, and 3) UPR profile of dermal fibroblasts isolated at puberty corresponds to multiplex stress resistance (ER stress, oxidative stress, DNA damage) of lung fibroblasts isolated at adulthood. We found that only tunicamycin induced BiP expression was repeatable in skin and lung fibroblasts. Tunicamycin induced expressions of BiP, GRP94, and CNX in skin fibroblasts predicted resistance of lung fibroblasts to tunicamycin, (but not thapsigargin and other inducers of lethal stress), which is indicative for the pro-survival role of UPR during stress. Tunicamycin induced BiP expression in skin and lung fibroblasts also predicted multiple cellular bioenergetics parameters in lung fibroblasts.

**Statements and Declarations:** No competing interests declared. This work was supported by National Science Foundation grants IOS1453784 and OIA1736150 to W.R.H., IOS1755670 to the PGSC, and a National Science Foundation EPSCoR pilot grant to K.N.Y. The funders did not have any input into the content of the manuscript nor require approval prior to submission.

## Introduction

Ecological and evolutionary physiologists have long been interested in uncovering the molecular and physiological mechanisms underlying inter- and intraspecific variation in energy metabolism (Friedman et al. 1992; Hammond et al. 2000; Killen et al. 2017; Yap et al. 2017, 2019), and life-history performance (Stearns 1992; Zera and Harshman 2001; Lailvaux, S. P., & Husak 2014) of animals. Variation in energy metabolism and performance have vast implications in life-history traits such as longevity and reproduction.

Recent efforts to elucidate the physiological mechanisms underpinning individual variation in life-history performance have largely focused on bioenergetics and oxidative stress (Dowling and Simmons 2009; Williams et al. 2010; Selman et al. 2012; Zhang and Hood 2016). While significant progress has been made regarding the role of oxidative stress and bioenergetics in life-history performance of animals, empirical evidence has been equivocal at best. For instance, some studies have found positive relationships between energy expenditure and longevity (Johnson et al. 1963; Lints 1989; Zhang et al. 2021a), some have found no relationships between energy expenditure and longevity (Robert et al. 2007; Selman et al. 2008; Vaanholt et al. 2009), whereas others have found negative relationships between energy expenditure and longevity (Stuart and Brown 2006; Munshi-South and Wilkinson 2010). Similarly, there does not seem to be a consensus regarding the relationships between oxidative stress and longevity of animals, with some studies showing negative correlations (Arking et al. 2000; Brown-Borg and Rakoczy 2003), while other studies showing no relationships between animals’ longevity and metrics of oxidative stress (Selman et al. 2008; Jang et al. 2009; Zhang et al. 2009). To some extent, these discrepancies in findings can be attributed to differences in study species and populations, as well as differences in research methodologies. However, the discrepancies could also be due to our inadvertent neglect of other physiological systems, such as ER stress (Chakrabarti et al. 2011; Pluquet et al. 2015; Yap et al. 2021) and DNA repair (Hart et al. 1979; Promislow 1994; Hyun et al. 2008).

One underappreciated area that could play a role in mediating variation in life-history performance is the unfolded protein response (UPR); a cellular stress response that during its adaptive phase reduces secretory protein load, enhances endoplasmic reticulum (ER) protein folding, and increases clearance capacity during homeostasis dysregulation and stress. More specifically, under unstressed conditions, properly folded proteins are released into the cytosol from the ER, and the ER chaperones binding immunoglobulin (BiP), calnexin (CNX), and glucose-regulated protein 94 (GRP94) are bound to receptors that prevent activation of the UPR. Under mild stress condition, where moderate amount of misfolded proteins are found in the ER, the chaperones dissociate from the receptors and induces activation of X-box binding protein-1 (XBP-1) and cleavage of Activating transcription factor 6 (ATF6), resulting in the adaptive phase of UPR where misfolded proteins are degraded. Under severe stress, where there is an overload of misfolded protein in the ER, the chaperone BiP dissociates from Protein Kinase R-like ER Kinase (PERK) to activate an apoptotic pathway via C/EBP homologous protein (CHOP) (Cao and Kaufman 2012; Bravo et al. 2013; Yap et al. 2021). ER stress and UPR have been studied extensively in the biomedical field, but they have received very little attention by ecological and evolutionary physiologists. ER stress and UPR play important roles in a suite of life-history traits, such as growth and development, physical performance, reproduction, and ageing (reviewed in Yap et al. 2021). Manipulation of ER stress is usually achieved via pharmacological means such as administration of tunicamycin and thapsigargin, which inhibits N-glycosylation and sarco/endoplasmic reticulum calcium ATPase, respectively (Yap et al. 2021). Furthermore, given that the ER and mitochondria interact to regulate cellular homeostasis (Lebiedzinska et al. 2009; Filadi et al. 2017; Carreras-Sureda et al. 2017), it seems intuitive that UPR phenotype would correlate strongly with mitochondrial physiology, which in turn would contribute to variations in whole-organism metabolism and life-history performance.

One of the most popular ways researchers have been studying cellular controls of life-history traits such as ageing and longevity is by assessing stress resistance and bioenergetic properties of primary dermal fibroblasts (Miller et al. 2011; Jimenez et al. 2018). Fibroblasts are one of the most abundant cell types in vertebrates and are involved in key biological processes like wound healing and connective tissue generation (Sorrell and Caplan 2004). It has been shown in many studies that primary fibroblasts isolated from long-lived animals have higher resistance to multiple forms of stress (Salmon et al. 2009; Harper et al. 2011; Miller et al. 2011; Jimenez et al. 2018). Primary dermal fibroblasts of long-lived naked mole rats (*Heterocephalus glaber)* also had lower rates of metabolism and can tolerate proteotoxic stress more effectively than mouse (*Mus musculus*) fibroblasts (Swovick et al. 2021). Additionally, a recent study by Havighorst et al. (2019) demonstrated that in genetically diverse deer mice (*Peromyscus maniculatus*), the UPR profile of fibroblasts isolated at puberty predicts physiological regulation of lipid metabolism and diet-induced hepatic steatosis later in life. Another recent study from the same group also found that the UPR profile of deer mice fibroblasts predicts skin inflammation and body weight gain after high-fat diet administration (Zhang et al. 2021b). These factors have been correlated with reduced physical performance and reduced lifespan in several other investigations (Johnson et al. 1997; Caruso et al. 2004; Martin et al. 2010).

However, it is unclear if findings from dermal fibroblasts can be generalized to other cell and tissue types, and if fibroblasts’ phenotypes (e.g., UPR profile) are repeatable across different life-history stages of an animal. To establish dermal fibroblasts as a reliable biomarker for studies of life-history performance, we need to first understand how physiological profiles of dermal fibroblasts match up with physiological profiles of fibroblasts isolated from other tissue sources such as lung fibroblasts, and how repeatable the physiological profiles are across time. Therefore, this study aimed to explore the relationships between UPR phenotype, cellular respiration, and stress resistance using primary dermal fibroblasts isolated at puberty and primary lung fibroblasts isolated at adulthood. Specifically, we are testing if 1) UPR profile of dermal fibroblasts isolated at puberty corresponds to UPR profile of lung fibroblasts isolated at adulthood, 2) UPR profile of dermal fibroblasts isolated at puberty and lung fibroblasts isolated at adulthood correspond to cellular bioenergetics of lung fibroblasts isolated at adulthood, and 3) UPR profile of dermal fibroblasts isolated at puberty corresponds to multiplex stress resistance (ER stress, oxidative stress, DNA damage) of lung fibroblasts isolated at adulthood.

## Materials and Methods

### Animals

Twenty-eight deer mice (*P. maniculatus*) were obtained from the Peromyscus Genetic Stock Center (PGSC), University of South Carolina (USC), Columbia, South Carolina. Dermal fibroblasts were isolated from ear punches collected at the time of weaning during standard marking procedure (more details below). All animals were transported to Auburn University and were allowed to acclimate to a semi-natural environment for 6 months. All husbandry and experimental procedures were approved by the Auburn University Institutional Animal Care and Use Committee (PRN 2019-3497). All animals were provided rodent chow (Teklad Global Diet 2019, Envigo, Indianapolis, IN) and water *ad libitum*. Animals were exposed to a natural light and dark cycle, as well as natural ambient temperatures and humidity.

### Dermal fibroblast culture and tunicamycin treatment

Dermal fibroblasts were isolated, cultured, and treated with tunicamycin following the protocol described in Havighorst et al. (2019). Briefly, ear punches collected at weaning were digested in collagenase I (at 100 U/ml) for 1 h to release fibroblasts from the tissue. Cells were cultured in RPMI-1640 + 10% fetal bovine serum + 500 U/ml penicillin + 500 μl/ml streptomycin+0.292 mg/ml L-glutamine in a humidified incubator with 5% CO_2_ in air. Cells were passaged when they were at 90% confluency or above. Cells were passaged for a maximum of 4 times before tunicamycin treatment. For tunicamycin treatment, dermal fibroblasts were cultured in 6-well plates and left for 24 h before addition of tunicamycin at 5µg/ml. Cells were exposed to tunicamycin for 5 h before RNA extraction.

### Lung fibroblast isolation and culture

Upon completion of the 6-month acclimation period at Auburn University, all animals were sacrificed by decapitation. Lung fibroblasts were isolated and cultured following Seluanov et al. (2010). Briefly, a small piece of lung tissue was excised immediately following decapitation, washed for 30 s in 70% EtOH and moved to DMEM medium. Lung tissue was digested in Liberase (at 100 U/ml) for 30 m to release fibroblasts from tissue. Cells were cultured in DMEM + 10% fetal bovine serum + 500 U/ml penicillin + 500 μl/ml streptomycin in a humidified incubator with 5% CO_2_ in air. Cells were passaged at 90% confluency. Upon passaging, cells were split into several culture dishes and were cultured in EMEM + 10% fetal bovine serum + 500 U/ml penicillin + 500 μl/ml streptomycin. Cells were passaged for a maximum of 3 times before tunicamycin treatment, cellular respiration measurements, and lethal stress assays. For tunicamycin treatment, lung fibroblasts were cultured in 6-well plates and left for 24 h before addition of tunicamycin at 5µg/ml. Cells were exposed to tunicamycin for 5 h before RNA extraction.

### RNA extraction, cDNA synthesis and qPCR

RNA extraction, complementary DNA (cDNA) synthesis, and quantitative PCR (qPCR) for fibroblasts were conducted following Havighorst et al. (2019) and Ruple et al. (2021). Oligonucleotide sequences (*BiP, GRP94, CNX, Gapdh*) used for qPCR amplification are shown in Table 1. Arbitrary units of target mRNA were normalized to the expression of *Gapdh*, and calculated using the 2^ΔΔ-Cq^ method where 2^Δ-Cq^ = 2^(housekeeping gene (HKG) Cq − gene of interest Cq) (Ruple et al. 2021). *BiP, GRP94, CNX* were chosen as our genes of interest because they are key chaperones that enhance protein folding capacity during the adaptive phase of UPR in the ER (Yap et al. 2021). They have been shown to be upregulated upon exposure to chemical inducers of ER stress and there is considerable individual variations regarding their degree of upregulations and expressions (Havighorst et al. 2019; Zhang et al. 2021b).

**Table 1:**
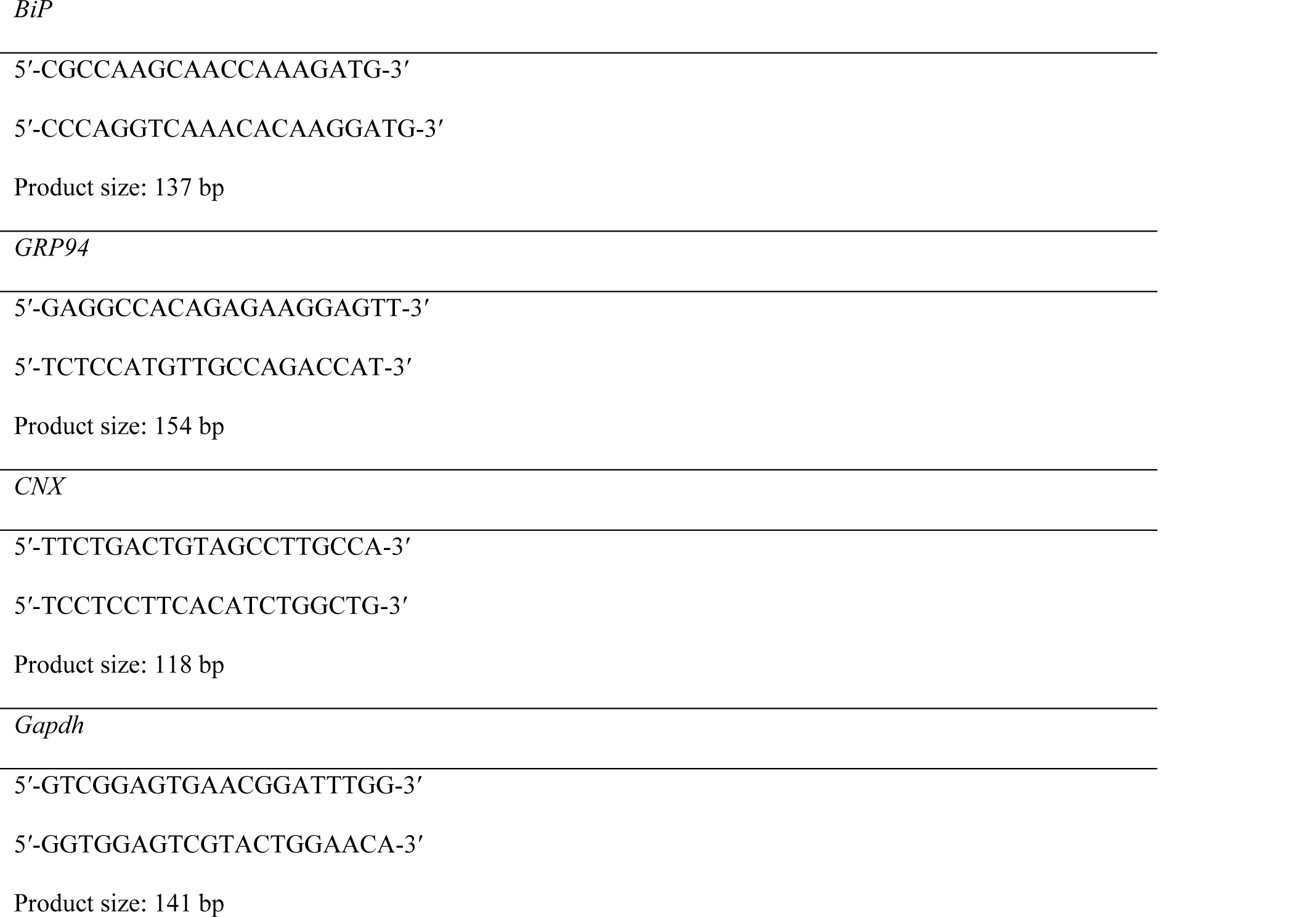
List of primers used.

### Assessment of lung fibroblast resistance to lethal stress

Assessments of cytotoxicity after exposure to lethal stress were conducted following Salmon et al. (2009) and Harper et al. (2011), with slight modifications. Lung fibroblasts (∼ 3 × 10^5^ cells) were seeded into a 96-well tissue culture-treated microtiter plate at a volume of 100 µl/well and were allowed to incubate for 24 h at 37°C in a humidified incubator with 5% CO_2_ in air. At the end of the incubation period, cells were washed with 1X PBS and incubated in serum-free EMEM for another 24 h. Thereafter, cells were exposed for 5 h to graded doses of one of tunicamycin, thapsigargin, paraquat, hydrogen peroxide (H_2_O_2_), and methylmethane sulfonate (MMS). Tunicamycin and thapsigargin are both ER stress inducers. Tunicamycin induces ER stress by inhibiting N-linked glycosylation, while thapsigargin induces ER stress by inhibiting Ca^2+^ ATPase of the ER. Paraquat and H_2_O_2_ are both potent oxidants that induce oxidative stress. MMS is a DNA alkylating agent that damages DNA by causing double-stranded breaks. Immediately following cytotoxic stress exposures, cells were washed once with 1X PBS, and incubated in serum-free EMEM for an additional 24 h. Cell survival was measured following the supplied protocol using EZViable™ Calcein AM Cell Viability Assay Kit (BioVision).

### Cellular respiration measurements

The rates of respiration (OCR) in lung fibroblasts were measured at 37°C using an oximeter (Oxytherm, Hansatech Instruments, UK). Cells were first suspended in culture media and respiration was measured for 5 m in the chamber. While fibroblasts are typically attached cells in culture, previous studies have shown that transient suspension of fibroblasts does not affect their respiratory activity and coupling control (Zdrazilova et al. 2022). This was immediately followed up with 1µM Oligomycin, ATP synthase inhibitor, added into the chamber for three-minutes. Then 1 µM FCCP, a mitochondrial uncoupler, was added and recorded for five-minutes. Finally, 0.5µM Rotenone, mitochondrial Complex I inhibitor, and 0.5µM Antimycin, Complex III inhibitor, were subsequently added and respiration was recorded for five to ten-minutes. Respiration rates were normalized to cell protein content. We measured basal respiration, proton leak, FCCP-induced maximal respiration, non-mitochondrial respiration, spare respiratory capacity, ATP production, and coupling efficiency following Winward, J. D., Ragan, C. M., & Jimenez (2018). Spare respiratory capacity, which indicates the cell’s capacity to respond to periods of high ATP demand, is calculated by subtracting basal respiration from FCCP-induced maximal respiration.

### Data analysis

Analyses were carried out using R version 0.99.467 (R Core Team 2013). Linear regressions were employed to test whether UPR profile of dermal fibroblasts isolated at puberty correspond to UPR profile of lung fibroblasts isolated at adulthood. Specifically, we analyzed how tunicamycin-induced gene expressions of BiP, GRP94, and CNX in dermal fibroblasts vary with tunicamycin-induced gene expressions of BiP, GRP94, and CNX in lung fibroblasts. The concentration or dose needed to kill 50% of the cells (LD_50_) was calculated using the FORECAST function in Excel for each of the lethal stressors (Harper et al. 2011). Cells were tested in triplicate, with one microtiter plate per stressor. We again employed linear regressions to investigate how tunicamycin-induced gene expressions of BiP, GRP94, and CNX in dermal fibroblasts vary with LD_50_ of tunicamycin, thapsigargin, paraquat, MMS, and H_2_O_2_ in lung fibroblasts. We then analyzed how tunicamycin-induced gene expressions of BiP, GRP94, and CNX in dermal fibroblasts vary with various OCR parameters in lung fibroblasts using linear regressions, with animals’ body mass as covariates. Sex was initially included in all models as either fixed effect, but was subsequently removed as it did not affect the analyses. Degrees of freedom, slope, intercept, R-squared values, and p-values were reported.

## Results

### UPR profile of dermal fibroblast and lung fibroblast

Tunicamycin induced BiP expression was positively correlated in dermal fibroblasts isolated at puberty and lung fibroblasts isolated at adulthood (*df* = 12, y = 0.69x – 2.55, p = 0.04, *R^2^* = 0.20, Fig. 1A). However, neither tunicamycin induced *GRP94* (*df* = 11, y = −0.42 x + 2.43, p = 0.19, *R^2^* = 0.07, Fig. 1B) nor *CNX* (*df* = 11, y = −0.53x + 1.23, p = 0.58, *R^2^* = 0.06, Fig. 1C) expressions were positively correlated in dermal fibroblasts isolated at puberty and lung fibroblasts isolated at adulthood.

**Figure 1:**
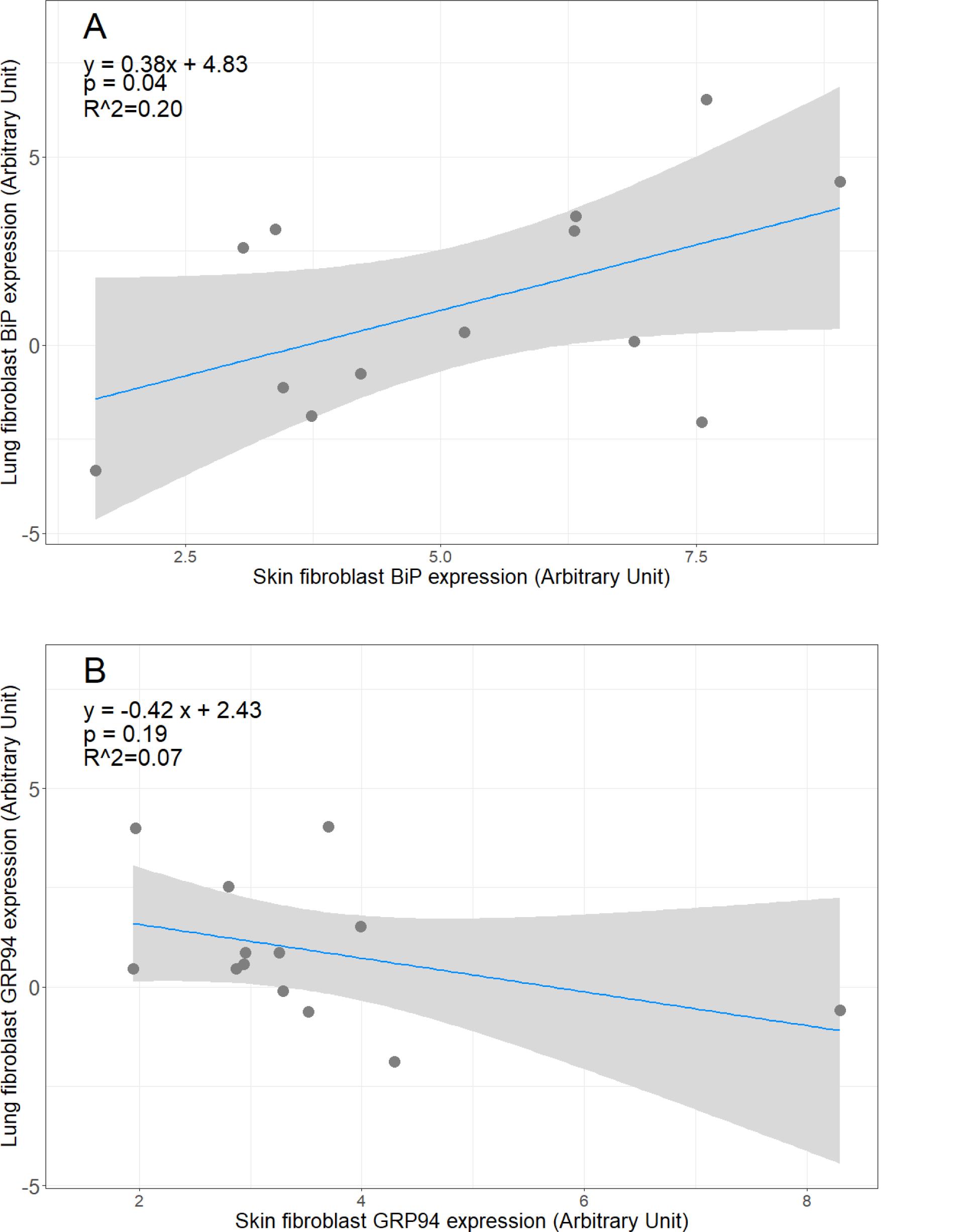
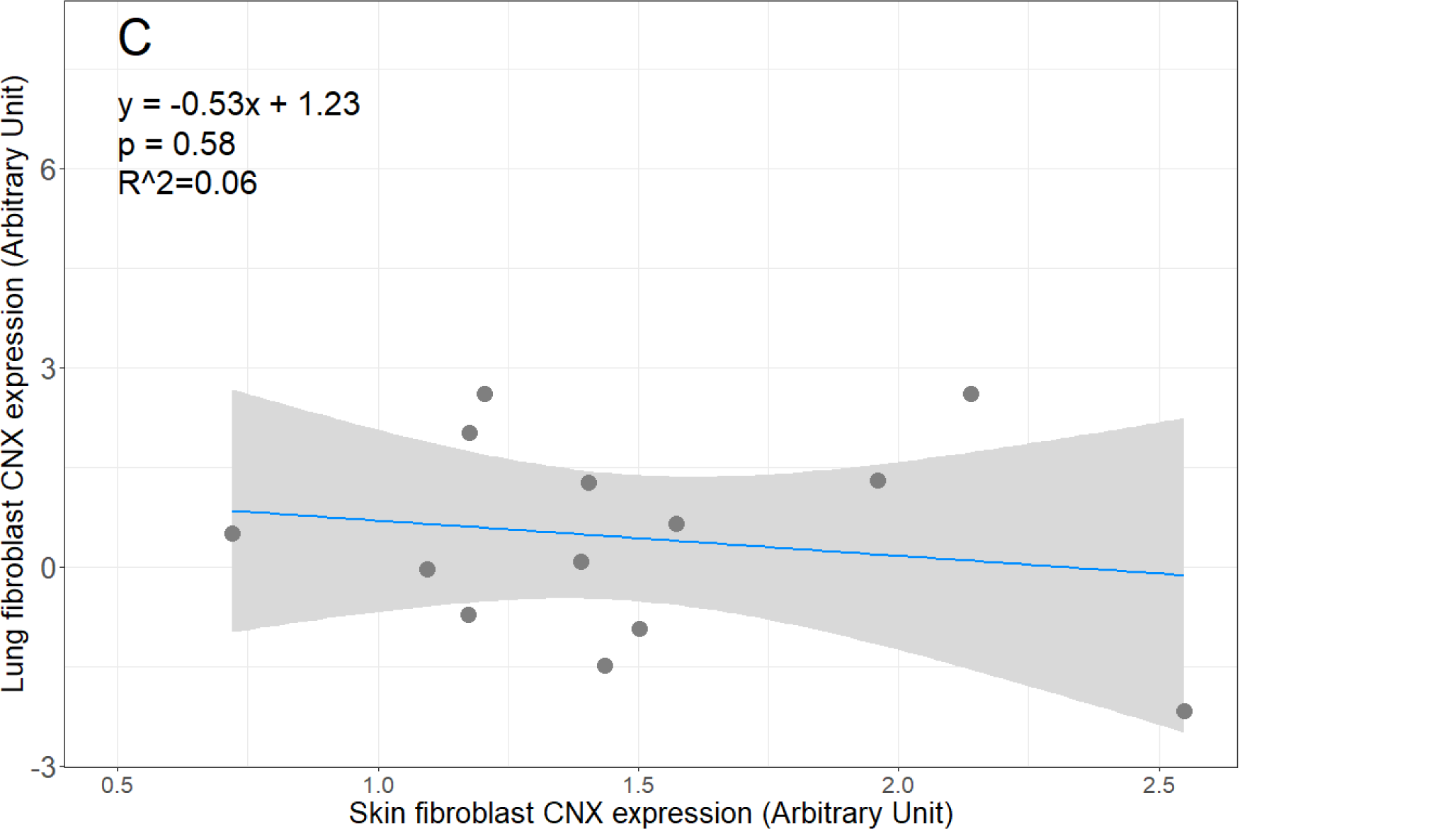
Relationships between A) tunicamycin induced BiP expressions in dermal fibroblasts and lung fibroblasts, B) tunicamycin induced GRP94 expressions in dermal fibroblasts and lung fibroblasts, and C) tunicamycin induced CNX expressions in dermal fibroblasts and lung fibroblasts.

### UPR profile of dermal fibroblast and resistance of lung fibroblasts to ER stress, oxidative stress, and DNA damage

Tunicamycin induced BiP expression (*df* = 15, y = 0.98x + 6.82, p < 0.01, *R^2^* = 0.54, Fig. 2A), GRP94 expression (*df* = 15, y = 1.28x + 8.15, p = 0.02, *R^2^* = 0.27, Fig. 2B), CNX expression (*df* = 15, y = 4.85x + 5.29, p < 0.01, *R^2^* = 0.34, Fig. 2C) in dermal fibroblasts isolated at puberty were all positively and significantly correlated with LD_50_ of tunicamycin in lung fibroblasts isolated at adulthood. The relationships were not significant once the sample with the highest BiP expression was removed from the analyses. However, the data point was kept in the analyses as it was not revealed to be a statistical outlier based on Grubbs test (p > 0.05 for all data points). None of the UPR markers were significantly related to LD_50_ of thapsigargin, paraquat, MMS, and H_2_O_2_ in lung fibroblasts isolated at adulthood (p > 0.10 in all cases).

**Figure 2:**
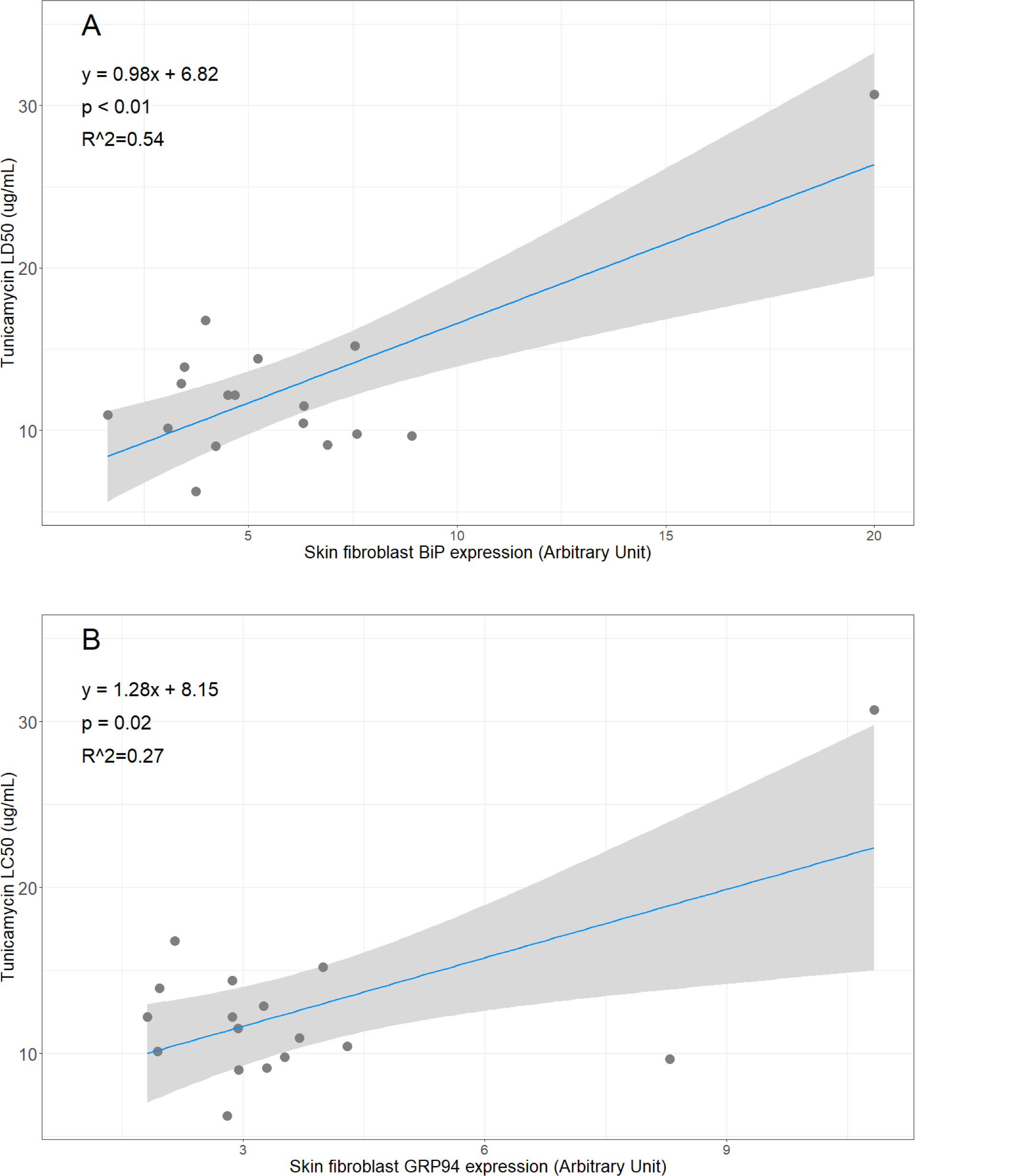
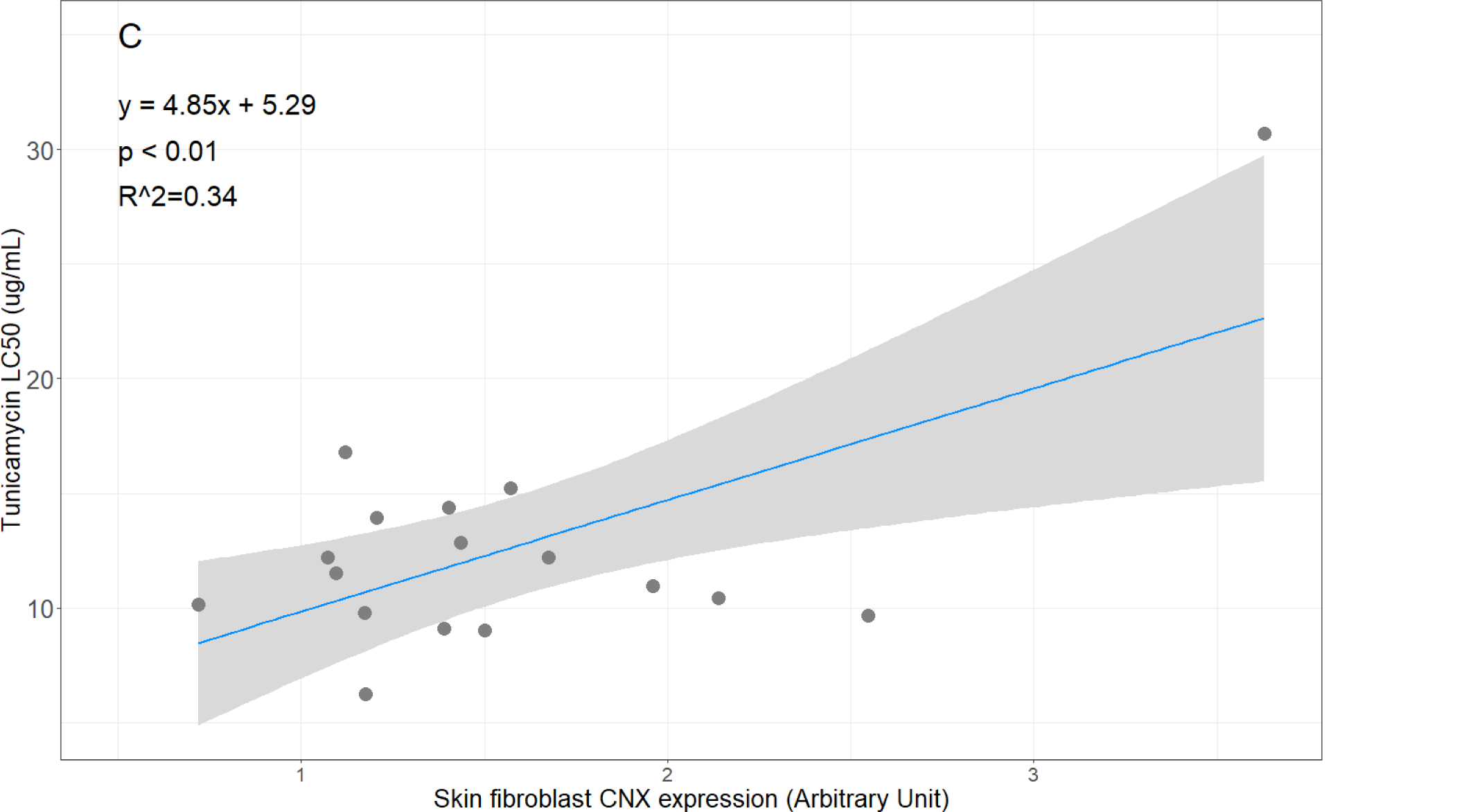
Relationships between A) tunicamycin induced BiP expressions in dermal fibroblasts and LD_50_ of tunicamycin in lung fibroblasts, B) tunicamycin induced GRP94 expressions in dermal fibroblasts and LD_50_ of tunicamycin in lung fibroblasts, and C) tunicamycin induced CNX expressions in dermal fibroblasts and LD_50_ of tunicamycin in lung fibroblasts.

### UPR profile of dermal and lung fibroblasts and cellular bioenergetics of lung fibroblasts

Tunicamycin induced BiP expression in dermal fibroblasts isolated at puberty was positively and significantly correlated with basal cellular respirations (*df* = 13, y = 6.11x + 157.37, p = 0.02, *R^2^* = 0.28, Fig. 3A) and proton leak (*df* = 13, y = 2.84x + 83.77, p < 0.01, *R^2^*= 0.48, Fig. 3B) in lung fibroblasts isolated at adulthood. Similarly, tunicamycin induced BiP expression in lung fibroblasts was positively and significantly correlated with basal cellular respirations (*df* = 15, y = 4.35x + 39.50, p < 0.01, *R^2^* = 0.33) and proton leak (*df* = 15, y = 1.44x + 26.95, p = 0.04, *R^2^* = 0.20) in lung fibroblasts. Similar positive relationship was found for tunicamycin induced BiP expression in dermal fibroblasts and FCCP-induced maximal respiration (*df* = 13, y = 12.88x + 258.40, p = 0.05, *R^2^* = 0.15, Fig. 3C), as well as tunicamycin induced BiP expression in lung fibroblasts and FCCP-induced maximal respiration (*df* = 15, y = 7.43x + 89.04, p = 0.08, *R^2^* = 0.14, but the relationship was only marginally significant. No significant relationships were found for tunicamycin induced BiP expression in dermal and lung fibroblasts and non-mitochondrial respiration, and spare respiratory capacity (p > 0.10 in all cases).

**Figure 3:**
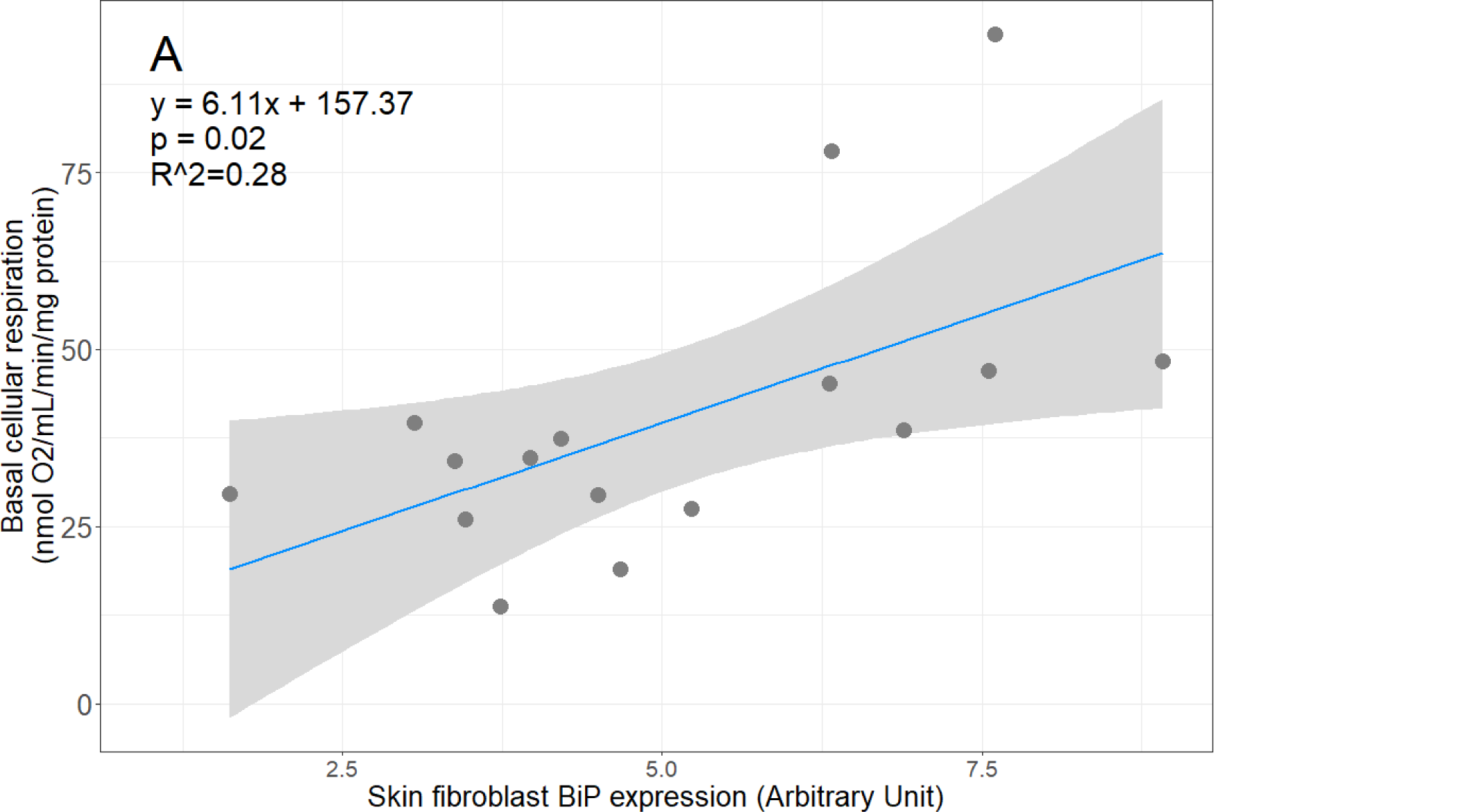
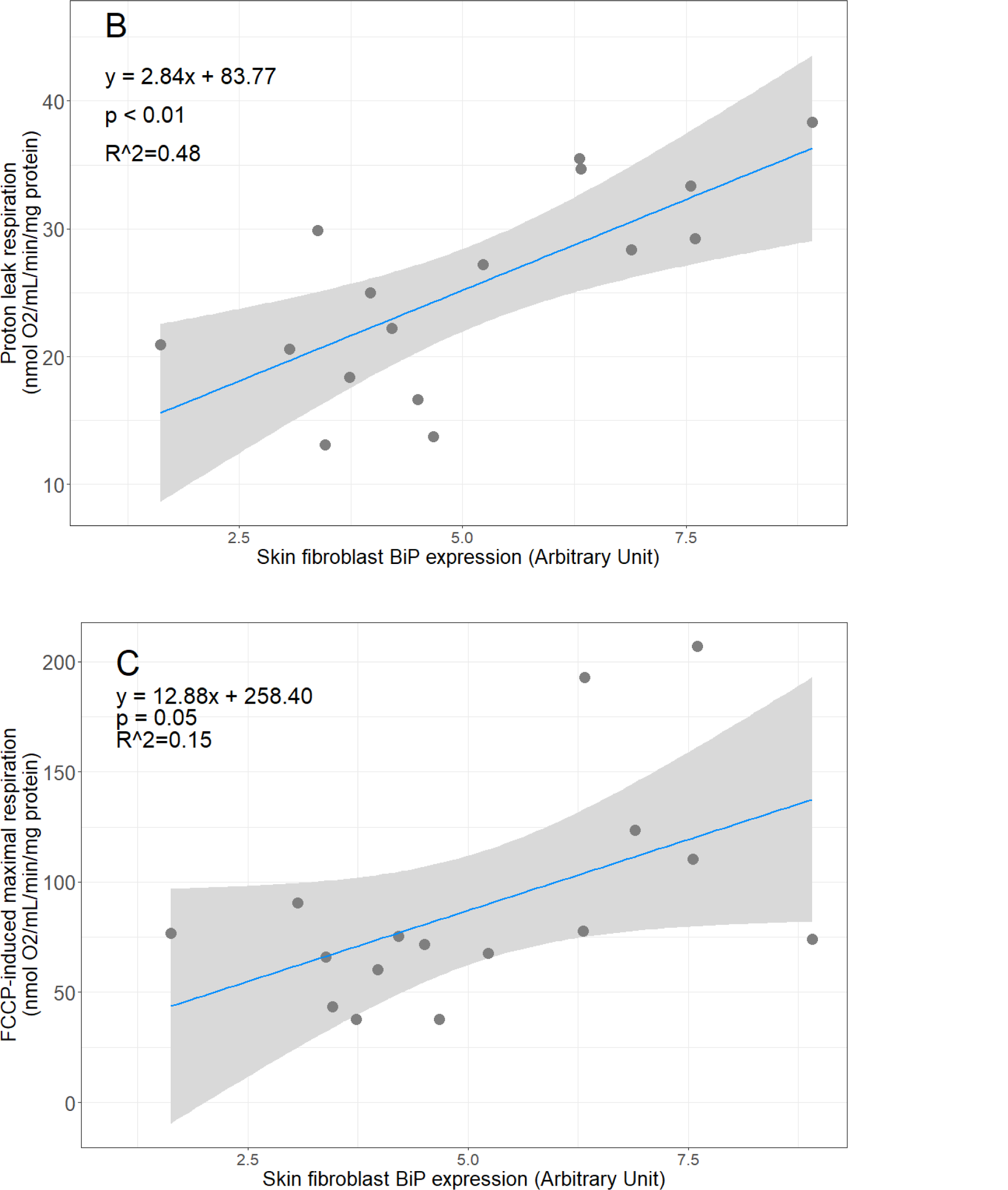
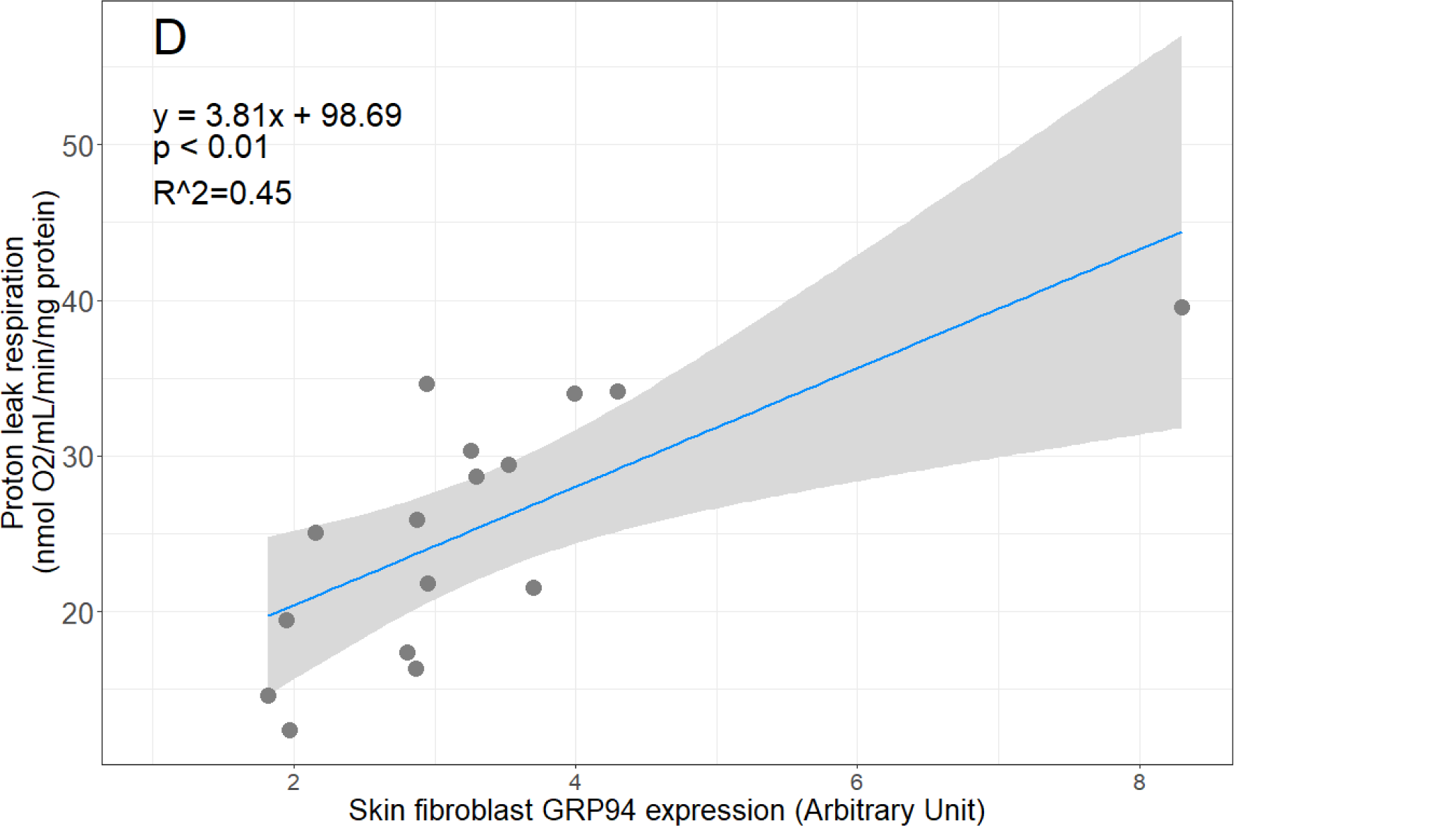
Relationships between A) tunicamycin induced BiP expressions in dermal fibroblasts and basal respiration in lung fibroblasts, B) tunicamycin induced BiP expressions in dermal fibroblasts and proton leak respiration in lung fibroblasts, C) tunicamycin induced BiP expressions in dermal fibroblasts and FCCP-induced maximal respiration in lung fibroblasts and D) tunicamycin induced GRP94 expressions in dermal fibroblasts and proton leak respiration in lung fibroblasts.

Tunicamycin induced GRP94 expression in dermal fibroblasts isolated at puberty were positively and significantly correlated with proton leak (*df* = 13, y = 3.81x + 98.69, p < 0.01, *R^2^* = 0.45, Fig. 3D) in lung fibroblasts isolated at adulthood. A negative relationship was found for tunicamycin induced GRP94 expression in lung fibroblasts and proton leak (*df* = 15, y = −2.25x + 30.60, p = 0.05, *R^2^* = 0.18), but the relationship was only marginally significant. All other cellular respiration metrics were not significantly related to tunicamycin induced GRP94 expression in dermal and lung fibroblasts (p > 0.10 in all cases). Similarly, tunicamycin induced CNX expression in dermal and fibroblasts isolated at puberty were not related to any cellular respiration metrics in lung fibroblasts isolated at adulthood (p > 0.10 in all cases).

## Discussion

This study aimed to compare physiological profiles of fibroblasts isolated from skin and lung tissues during puberty and adulthood respectively. Specifically, we investigated the relationship between UPR phenotype, cellular respiration, and stress resistance using primary dermal fibroblasts isolated at puberty and primary lung fibroblasts isolated at adulthood. In terms of repeatability of UPR profile, we found that only tunicamycin induced BiP expression was repeatable in skin and lung fibroblasts. Regarding resistance to lethal cytotoxic stress, tunicamycin induced expressions of BiP, GRP94, and CNX in skin fibroblasts predicted resistance of lung fibroblasts to tunicamycin, but not thapsigargin and other inducers of lethal stress. Additionally, tunicamycin induced BiP expression in skin fibroblasts also predicted multiple cellular bioenergetics parameters in lung fibroblasts.

Tunicamycin induced BiP, but not GRP94 and CNX expressions showed moderate repeatability in skin and lung fibroblasts. This suggests that although some aspects of UPR profile are consistent across fibroblasts isolated from different tissue sources and across different life-history stages, this is likely to be chaperone-dependent. Many previous studies have often assumed that regardless of tissue origin, fibroblasts are uniform cell population with comparable functions. However, more and more studies have shown that this is not necessarily true (Fries et al. 1994; Phipps et al. 1997; Lindner et al. 2012). For instance, Lindner et al. (2012) showed that heart, skin, and lung fibroblasts had very different basal expressions of various metalloproteases, and they displayed different responses to TNF-α exposure. It is unclear why we observed repeatability of BiP expression, but not GRP94 and CNX in skin and lung fibroblasts, although it has been shown previously that N-linked glycosylation does not have any known effect on GRP94 activity (Marzec et al. 2012). The major chaperone BiP is considered the central regulator of ER stress as it controls the activation of ER stress sensors such as IRE1, PERK, and ATF6 (Lee 2005; Ni and Lee 2007; Yap et al. 2021). GRP94 is a HSP90-like protein that also functions as a chaperone but unlike BiP, has more substrate specificity (Ni and Lee 2007; Marzec et al. 2012). Similarly, compared to BiP, CNX has more substrate specificity as it is responsible for folding of glycoproteins (Ni and Lee 2007). It is also possible that the functions and specificity of the chaperones are slightly different in different tissues, but we are not aware of any studies that have explored this.

However, despite not observing strong repeatability in tunicamycin induced expressions of GRP94 and CNX in skin and lung fibroblasts, all three UPR markers predicted resistance of lung fibroblasts to tunicamycin. This observation could be attributed to the positive correlations observed between the expressions of BiP, GRP94, and CNX. Although both tunicamycin and thapsigargin are both ER stress inducers, the expressions of BiP, GRP94, and CNX failed to predict the cells resistance to thapsigargin in lung fibroblasts. The doses of tunicamycin and thapsigargin used in this study induced cell death at similar levels and therefore are comparable. The fact that expressions of BiP, GRP94, and CNX failed to predict the cells resistance to thapsigargin in lung fibroblasts is perhaps not surprising considering tunicamycin and thapsigargin induce ER stress via different mechanisms, suggesting that the type of ER stress, or how ER stress is induced, needs to be taken into consideration when studying physiological effects of ER stress. Indeed, other studies have shown differential physiological effects of tunicamycin-induced and thapsigargin-induced ER stress as well (Sasaya et al. 2008; Brodnanova et al. 2021). Tunicamycin induced expressions of all UPR markers in skin fibroblasts also did not predict resistance of lung fibroblasts to oxidative stress and DNA damage. This pattern of differential resistance suggests that cells’ responses to various forms of cytotoxic stress are regulated and coordinated differently. This finding corroborates with findings from other studies looking at cellular stress resistance and aging, where studies often find cells from longer-lived animals more resistant to one form of cellular stress but not another (Salmon et al. 2009; Harper et al. 2011; Miller et al. 2011; Winward, J. D., Ragan, C. M., & Jimenez 2018).

Tunicamycin induced BiP expression and to some extent GRP94 expression of dermal fibroblasts isolated at puberty and lung fibroblasts isolated at adulthood predicted cellular bioenergetics parameters such as basal respiration, proton leak, and maximal respiration in lung fibroblasts isolated at adulthood. This suggests that UPR profile, or ER physiology in general, plays an important role in regulating bio-energetic profile of cells, either directly or indirectly. Indeed, many studies have shown that there is tight physical linkage between the ER and mitochondria and that the organelles interact closely to regulate cellular homeostasis (Giorgi et al. 2009; Bravo et al. 2011; Filadi et al. 2017; van Vliet and Agostinis 2018; Gordaliza-Alaguero et al. 2019). It should be noted that the observed relationship is correlational in nature. It remains to be determined whether induction or alleviation of ER stress affect bio-energetic capacity of fibroblasts from other tissue origins. It also remains to be seen whether animals with varying tolerance and resistance to ER stress have different bio-energetic profile.

Taken together, our findings indicate that some aspects of ER physiology and UPR profile are repeatable across fibroblasts isolated from different tissue type and across different life-history stages of an animal, but other aspects are regulated and expressed differently. Our findings also showed that not all forms of ER stress are created equal. Different forms of induction can elicit different molecular and physiological responses. Researchers studying ER stress should to be mindful of this when studying the physiological and functional implications of ER stress induction.

There are a few caveats in this study. First, it would be more informative if we could compare dermal fibroblasts from pubertal deer mice, dermal fibroblasts from adult deer mice, and lung fibroblasts from adult deer mice. However, we were unable to isolate and culture dermal fibroblasts from adult deer mice due to fungal contamination issues. The animals were housed in outdoor seminatural condition during the summer months in Auburn, Alabama, where heat and humidity resulted in fungal growth on hair and skin of animals. However, the animals appeared healthy throughout the study and no fungal contamination was detected in primary lung fibroblasts culture. It is also possible that the results we observed in this study were due to developmental or environmental effects. Furthermore, although the same dose (5µg/ml) was used for tunicamycin-induced gene expressions of dermal and lung fibroblasts, it should be noted that we did not establish dose titration curves on BiP expression for lung and dermal fibroblasts. Though unlikely, it is possible that a different dose of tunicamycin would elicit different responses in lung and dermal fibroblasts.

Dermal fibroblast is relatively easy to isolate and culture, and the procedure only requires a small skin biopsy and does not necessitate euthanasia of animals. Thus, more and more studies are using fibroblasts to characterize various physiological modalities of animals and to infer whole-organism phenotype (Williams et al. 2010; Miller et al. 2011; Jimenez et al. 2018; Zhang et al. 2021b). Our findings suggest that some aspects of ER physiology observed in dermal fibroblasts could be extrapolated to other cell types, or at least fibroblasts isolated from a different tissue origin. However, while it is promising that some aspects of ER physiology are repeatable, we need to be careful not to infer causation from correlational findings. We also need to be cautious about extrapolating all findings from primary dermal fibroblast to other cell types and to the whole-organism level. Future studies should strive to characterize UPR profile and ER physiology in other cell types and investigate whether and to what extent UPR profile is correlated in different cell and tissue types.

## Acknowledgement

We thank MaKalea Kirkland, LouAnn Crosby, Emma Rhodes, and Aubrey Taylor for their invaluable assistance with animal husbandry and laboratory assays. We also thank Shelby Osburn, Joshua Godwin, and Paulo Mesquita for technical assistance and general discussions in the laboratory.

